# Targeted Eicosanoid Profiling Reveals MicroRNA-155 Shifts PGE2/PGD2 Balance Towards Oncogenic State

**DOI:** 10.1101/2020.04.08.032136

**Authors:** Sinae Kim, Eun Sung Lee, Eun ji Lee, Jae Yun Jung, Sae Byul Lee, Hee Jin Lee, Jisun Kim, Hee Jung Kim, Jong Won Lee, Byung Ho Son, Gyungyub Gong, Sei Hyun Ahn, Suhwan Chang

## Abstract

microRNA-155 is a strong oncogenic microRNAs with multiple functions. To reveal its novel function in cancer metabolism, we performed a targeted metabolomic analysis of icosanoids, the key metabolites of inflammation-related carcinogenesis. We found miR-155-depleted cells expressed unbalanced prostaglandins, especially in PGE2 and PGD2. Subsequent analysis of primary cancer cells and 20 TNBC specimen indicated a positive correlation between miR-155 and PGE2/PGD2 ratio. Mechanistically, miR-155 controls the prostaglandin shift by up-regulating PGE2-producing enzymes PTGES/PTGES2 and down-regulating PGD2-producing enzyme PTGDS. Further analysis showed that the up-regulation of *PTGES2* is driven by miR-155-cMYC axis whereas *PTGES* is transactivated by miR-155-KLF4 axis, indicating a dual-regulatory mode for the metabolic enzyme expression drives a shift in PGE2/PGD2 balance. Lastly, we show the miR-155-driven cellular proliferation is significantly restored by the siRNA of PTGES1/2, of which expression also correlates with patients’ survival. Taken altogether, our study reveals an unprecedented oncogenic function of miR-155 on prostaglandin regulation.

## Introduction

microRNA-155 is encoded by a non-coding, BIC (B-cell lymphoma Insertion Cluster) gene, discovered as an oncogene that is activated by retroviral insertion (Eis et al, 2005). As miR-155 is the only microRNA produced from the BIC, the oncogenic function of the BIC gene is driven by miR-155. In addition to its function in lymphoid cell transformation (Kluiver et al, 2005; Tili et al, 2009), miR-155 has shown oncogenicity in solid tumors including lung, colon, pancreatic, and breast cancer (Jurkovicova et al, 2014). Based on the nature of incomplete miRNA-target base-paring, the number of miR-155 targets was estimated to be more than 200, which was verified by the Ago-CLIP results (Meier et al, 2013). Because of the large number of targets, it is difficult to dissect the molecular mechanism of the miR-155-induced oncogenicity. In cancer cells, miR-155 affects multiple biochemical, physiological aspects of cancer such as proliferation, migration, drug resistance, autophagy, and immune evasion (Faraoni et al, 2009; Lind & Ohashi, 2014; Wang et al, 2018). Notably, several recent reports revealed the function of miR-155 in cancer metabolism including glucose (Han et al, 2015; Jiang et al, 2012), thiamine(Kim et al, 2015) and estrogen metabolism(Bacci et al, 2016), thereby suggested the role of miR-155 on deregulated metabolic control in cancer cells.

Eicosanoids are 20-carbon bioactive lipids generated from fatty acids, with diverse biological functions related to cancer, including cell growth, migration, and angiogenesis (Hu et al, 2018; Wang & Dubois, 2010). The biogenesis of eicosanoids starts from arachidonic acid, generated from phospholipid by PLA2; subsequently, it is metabolized into leukotrienes and prostanoids (Serhan et al, 1996). Prostaglandin is a set of lipid metabolite of prostanoids with diverse physiological functions (Hata & Breyer, 2004), which are derived from arachidonic acids and have 20 carbons with a 5-carbon ring. For each of the prostaglandin, there is a synthase responsible for its biogenesis (Helliwell et al, 2004). Among the prostaglandins, PGE2 is well-known for its oncogenic function via the EP2 receptor followed by the PKA-CREB pathway, Ras-Raf-MAPK, or PI3K-AKT pathway (Nakanishi & Rosenberg, 2013). In contrast, PGD2, an antitumor prostaglandin, reduces the expression of cMYC and Cyclin D1 and increases apoptosis (Okuda-Ashitaka et al, 1990). Multiple studies have shown that microRNAs control the synthesis of eicosanoids: for example, Cox2, a key enzyme for the synthesis of PGH2 from arachidonic acid, is regulated by miR-101a, miR-146a, miR-16, and miR-26b (Hao et al, 2011; Kwon et al, 2015; Pham et al, 2013; Young et al, 2012). Similarly, 5-LO and FLAP, which are responsible for 5-HPETE biogenesis, are controlled by miR-219 and miR-135a, respectively(Lutz & Cornett, 2013). Despite these results, it is unclear how miR-155 contributes to eicosanoid homeostasis in cancer. In this study, we performed eicosanoids profiling in miR-155-deficient breast cancer cells and investigated the oncogenic role of miR-155 on prostaglandin biogenesis and its clinical impact on breast cancer.

## Results

### Alteration of prostaglandin metabolism in miR-155-deficient breast cancer cells

We have previously reported the role of miR-155 on the MDSC recruitment regulation (Kim et al, 2016) and glucose metabolism(Kim et al, 2018) by using a breast cancer mouse model with miR-155 deficiency. Conversely, another group showed that miR-155 enhances COX-2 expression and PGE2 secretion in asthmatic and nonasthmatic hASMC(Comer et al, 2015), raising the possibility that miR-155 has a role on prostaglandin expression in tumors as well. Therefore, we conducted a targeted metabolite profiling of eicosanoids, of which prostaglandins are a subfamily. As a model, we used miR-155-high mammary tumor and its isogenic, miR-155-depleted tumor generated by stable inhibition of miR-155. Our targeted LC-MS/MS analysis detected 30 isocyanides, including leukotrienes, thromboxanes, and prostaglandins. Fig 1A shows the decrease in PGE2/PGD2 ratio upon knockdown of miR-155 (raw data in Figure EV1 and Appendix Table S1). Using miR-155 ^KO/+^ and miR-155 ^KO/KO^ breast tumor models (Kim et al., 2016), we then examined if the miR-155 status correlates with PGE2/PGD2 expression in the plasma of the tumor model. Mice bearing miR-155 ^KO/KO^ tumors had less plasma PGE2 and more PGD2 than those bearing miR-155 ^KO/+^ tumors(Fig 1B, 1C and Appendix Table S2), consistent with the data in Fig1A.

**Figure 1.**
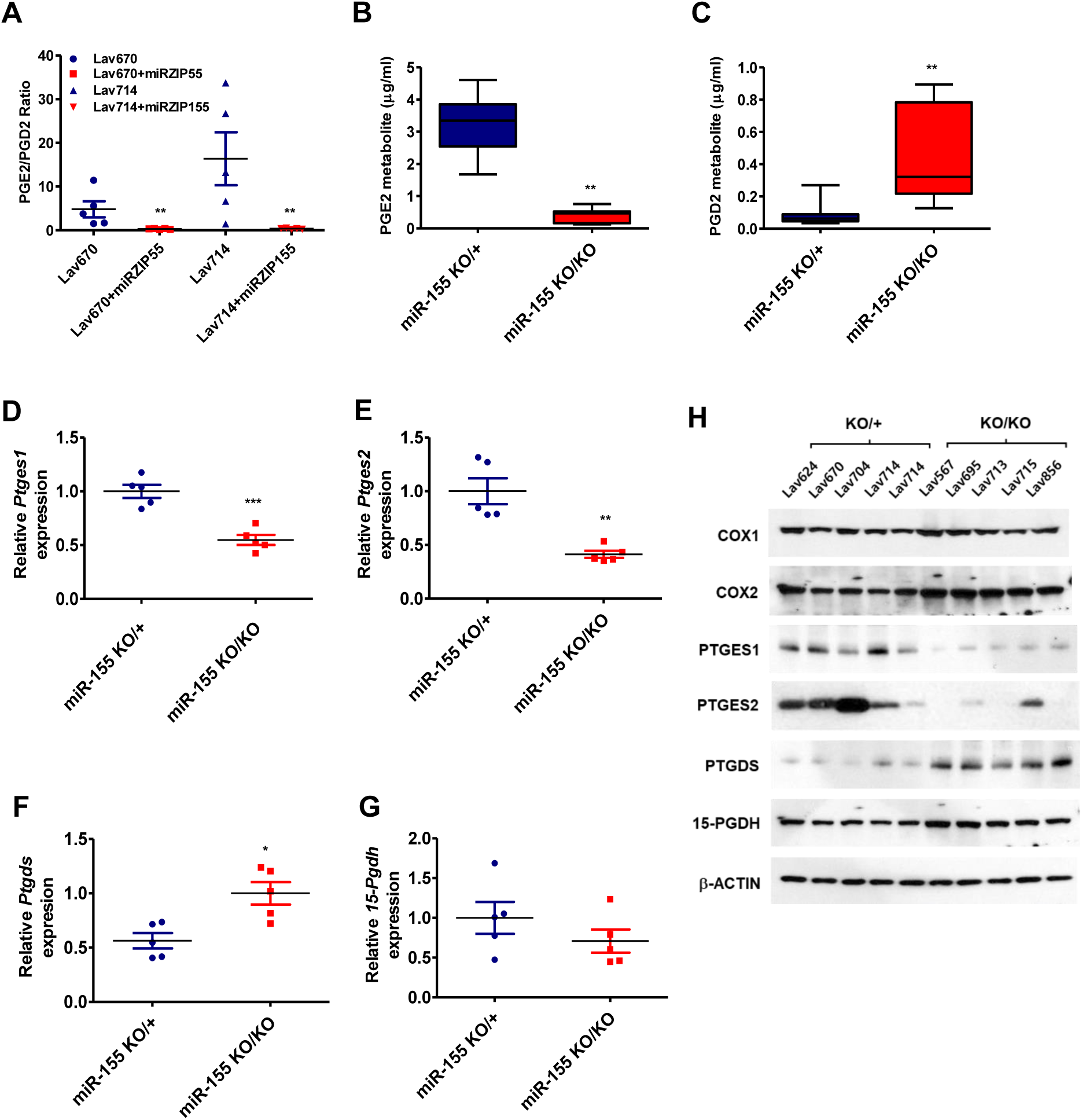
miR-155 depletion in murine mammary tumors reduces PGE2/PGD2 ratio and alters the expressions of related metabolic enzymes. A. PGE2/PGD2 ratio in miR-155-positive (Lav670 and Lav714) and miR-155-depleted (Lav670+miRZIP155 and Lav714+miRZIP155) tumors (n=20). B-C. PGE2(B) and PGD2(C) level in miR-155 ^KO/KO^ and in miR-155^KO/+^ tumors measured by ELISA (n=10). D-G. Relative RNA expression of *Ptges1*(D), *Ptges2*(E), *Ptgds*(F) and *15-Pgdh*(G) in miR-155 ^KO/KO^ and miR-155^KO/+^ tumors (n=10). H. Western blot analysis of key metabolic enzymes of PGE2/PGD2 production in miR-155 ^KO/KO^ and miR-155^KO/+^ tumors (n=10). COX1, COX2, PTGES1, PTGES2, PTGDS, and 15-PGDH expressions are shown. β-actin antibody was used to ensure equal loading.

PGE2 is known as an oncogenic metabolite, whereas PGD2 is a tumor suppressive metabolite. Therefore, the increase ratio of PGE2/PGD2 in miR-155-positive tumor and plasma may be a signature of oncogenic prostaglandin shift. We thus sought to delineate the target genes of miR-155 in association with the prostaglandin shift. There are three enzymes that control the level of PGE2: PTGES1 (Prostaglandin E2 Synthase 1), PTGES2, and 15-PGDH (15-Hydroxyprostaglandin dehydrogenase). The former two enzymes synthesize PGE2 from PGH2 (Nuttinck et al, 2008), and the 15-PGDH inactivates PGE2 by oxidizing the 15-hydroxyl group(Tai et al, 2002). Consistent with the positive correlation between microRNA-155 and the PGE2 level, we observed that miR-155-deficient tumors had decreased RNA levels of PTGES1 or PTGES2 (Fig 1D and 1E) and increased level of PTGDS (Fig 1F), while that of 15-PGDH was not significantly changed (Fig 1G). Western blot analysis in Fig 1H confirmed the RT-PCR data, showing down-regulated PTGES1/2 and up-regulated PTGDS in miR-155-deficient tumors. These data collectively suggest a strong interaction between miR-155 and prostaglandin expression, which is likely mediated by the regulation of enzymes responsible for the synthesis of PGE2/PGD2.

### microRNA-155 changes PGE2/PGD2 levels in human breast cancer cells via the regulation of PTGES/PTGES2/PTGDS expression

Based on the results obtained from the miR-155-deficient mouse model, we further examined the role of miR-155 on prostaglandin metabolism in human breast cancer cells. As our previous study revealed that miR-155 has a high expression level in MDA-MB-436 and Hs-578T cells and low expression level in MCF7 (Kim et al., 2018), we depleted miR-155 in MDA-MB-436 and Hs-578T cells and overexpressed miR-155 in MCF7 cells by lentivirus-mediated antagomir/mimic of miR-155, the results of which are shown in Figure EV 2A-2B. Consistent with the data from miR-155 KO mouse model, we observed decreased PGE2 and increased PGD2 levels in MDA-MB-436 and Hs-578T upon miR-155 depletion (Fig 2A, Figure EV 2C and Appendix Table S3 for raw data). Conversely, we observed increased PGE2 and decreased PGD2 in MCF7 cells upon miR-155 overexpression (Fig 2B). We further observed reduced PTGES1/2 and increased PTGDS expression levels in miR-155-depleted cells (Fig 2C-2E, red bars). In contrast, the overexpression of miR-155 in MCF7 cells resulted in increased PTGES1/2 and reduced PTGDS expression levels (Fig 2C-2E, yellow bars). We found no significant changes in 15-PDGH in all three cells tested (Fig 2F). These results were validated on the protein level by Western blot (Fig 2G).

**Figure 2.**
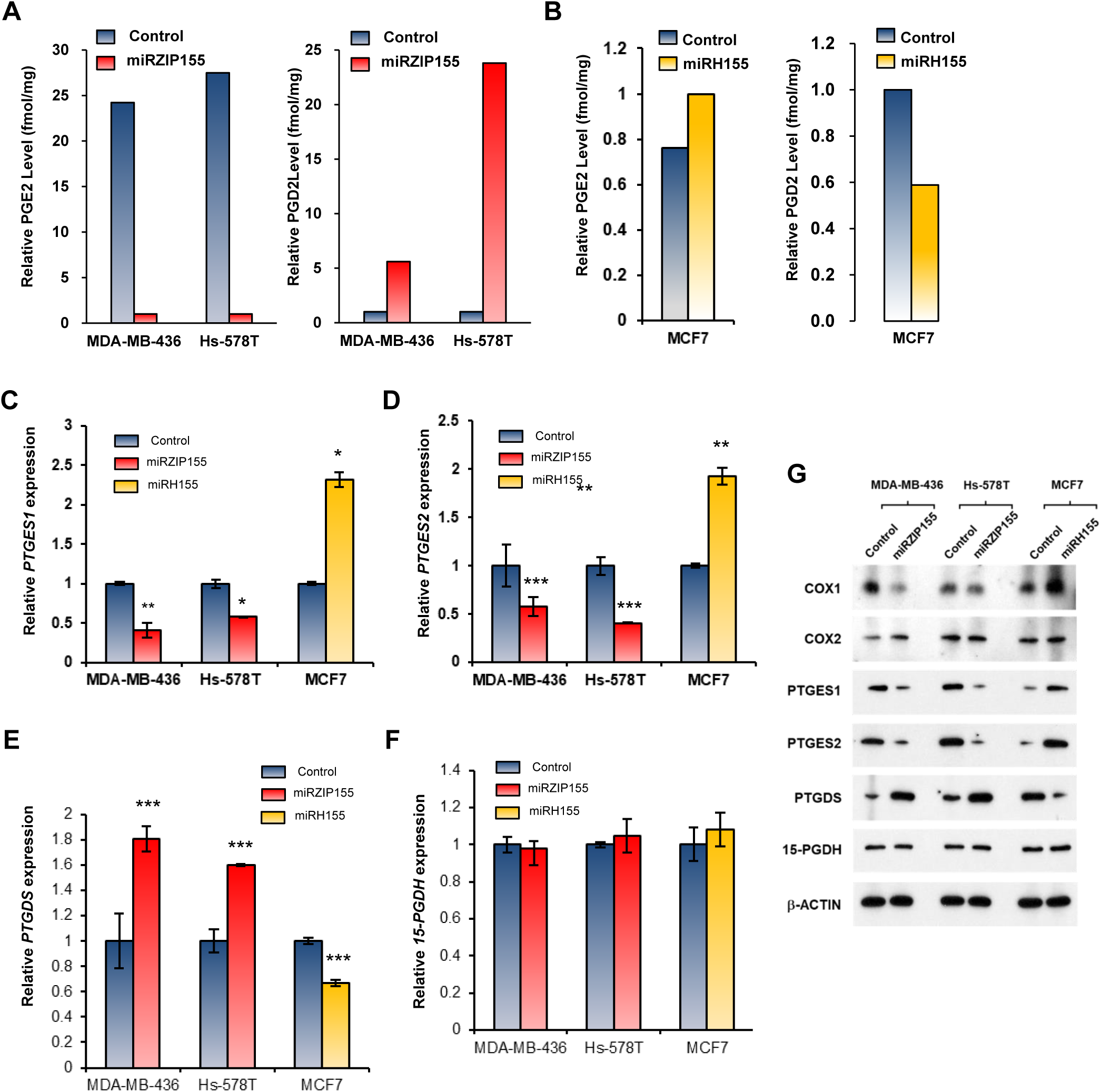
Modulation of miR-155 results in altered PGE2 and PGD2 production in human breast cancer cells, by the regulation of three prostaglandin synthases. A. Relative PGE2 (left) and PGD2 (right) levels in MDA-MB-436 and Hs-578T cells in control (in blue) or in miR-155-knockdown cells (miRZIP155 in red). B. Relative PGE2 (left) or PGD2 (right) level in MCF7 cells with overexpression of miR-155 (miRH, in yellow). C-F. Relative RNA expression of *Ptges1*(C), *Ptges2*(D), Ptgds(E) and *15-Pgdh*(F) in MDA-MB-436, Hs-578T, and MCF7 cells after knockdown (miRZIP155, in red) or overexpression (miRH155, in yellow) of miR-155. G. Western blot analysis of key metabolic enzymes for PGE2/PGD2 production in three breast cancer cell lines with after knockdown (miRZIP155) or overexpression (miRH155) of miR-155. β-actin antibody was used to ensure equal loading

We further confirmed our findings in PDCs and observed that the overexpression of miR-155 in three independent miR-155-low PDCs (PDCL1-3 in Fig 3A) replicated the results obtained in the miR-155-overexpressed MCF7 cell line in Fig 2 (Fig 3B-3F, raw data in Appendix Table S4). Importantly, miR-155 expression in PDCs restored the expression levels of PTGES1/2 and PTGDS to those of miR-155-high PDCs, and those results were validated by Western blot analysis (Fig 3G). Two of the miR-155-overexpressed PDCL (PDCL1 and 3) showed increased RNA levels of 15-PGDH (Fig 3H) but no significant changes in the protein level thereof (Fig 3G).

**Figure 3.**
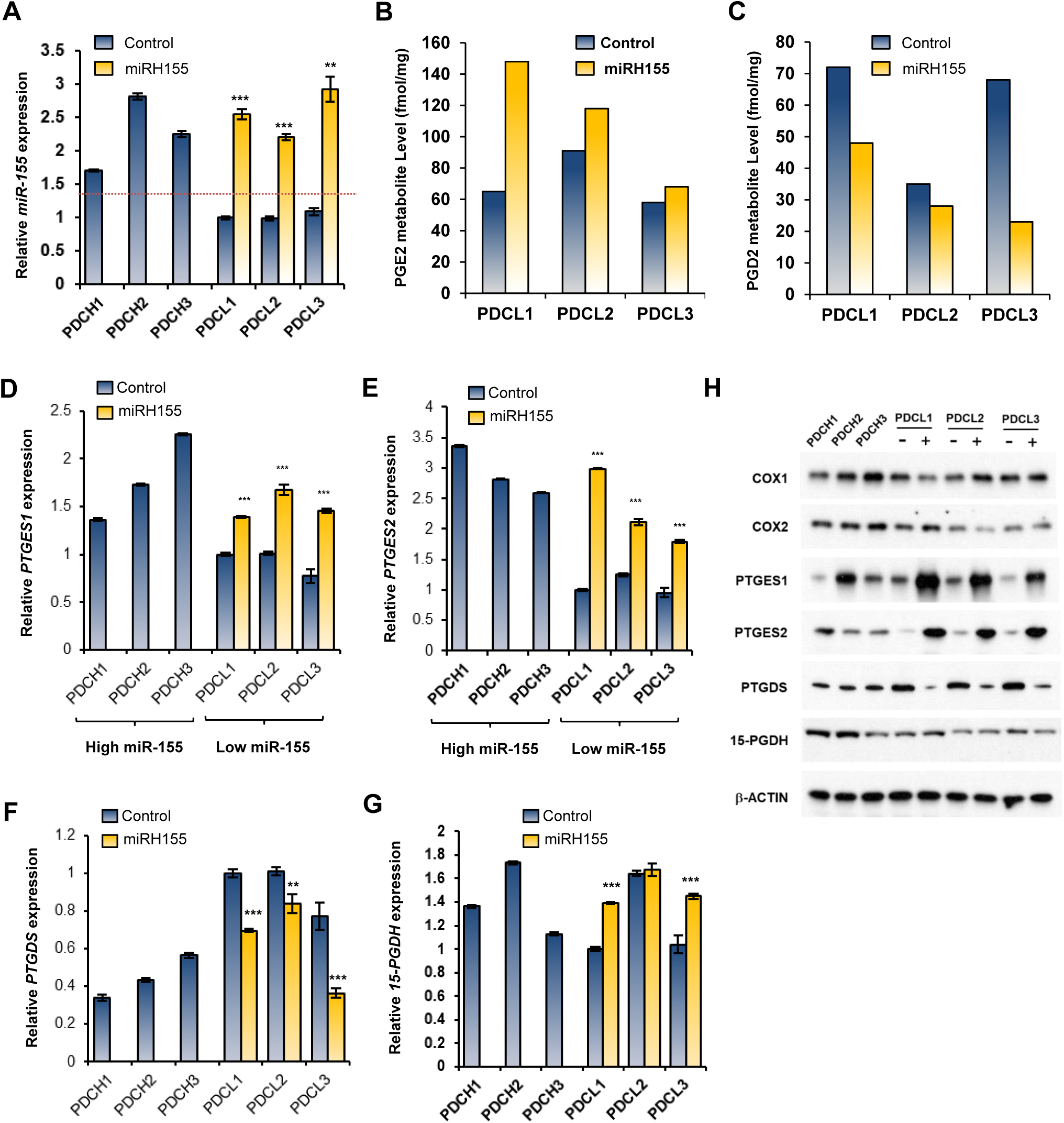
miR-155 overexpression shifts PGE2 and PGD2 production in patient-derived primary cancer cells (PDCs). A. Relative level of miR-155 in six PDCs. Three high-expressers were labeled as PDCH and low-expressers were labeled as PDCL. Yellow bars on PDCL (miRH 155) indicates miR-155 level expressed by lentiviral infection. B and C. The level of PGE2(B) or PGD2(C) in three PDCLs with overexpressed miR-155 (miRH155, in yellow). D-G. Relative RNA expression of *Ptges1*(D), *Ptges2*(E), *Ptgds*(F) and *15-Pgdh*(G) in PDCH, PDCL, and PDCL with lentiviral miR-155 expression (miRH155). H. Western blot analysis of key metabolic enzymes for PGE2/PGD2 production in three PDCH, PDCL, and PDCL with miR-155 expression (marked as +). β-actin antibody was used to ensure equal loading.

### PTGES2 expression is positively regulated by the miR-155-Myc axis

miR-155 regulates the expression of multiple genes including CEBP/beta, cMYC, FOXO3a, thiamine synthase, and GLUT1 via the direct regulation on 3’UTR or indirect regulation via the transcriptional regulator of the target genes(Kim et al., 2018; Kim et al., 2015; Kim et al., 2016). As we observed that the alteration of miR-155 level changes the expression levels of PTGES1/2 and PTGDS, we investigated if such regulation is carried out in a direct manner or via other transcription factors. Considering that miRNA mostly represses gene expression, positive correlation between PTGES1 or PTGES2 and miR-155 suggested an indirect regulation. Indeed, UTR analysis of the two genes in TargetScan revealed no putative miR-155 binding sites (Figure EV3). Therefore, we hypothesized a presence of mediators (possibly transcription factor) which are regulated by miR-155, is responsible for the control of PTGES1/2 gene expression. Fig 4A and 4B show that miR-155 knockdown reduces the promoter activity of PTGES2, suggesting that miR-155 regulates PTGES2 expression via promoter regulation. Therefore, we searched for transcription factor mediating this regulation, and among the candidates, we focused on cMYC as our previous study identified cMYC as a gene with a positive correlation with miR-155 (Kim et al., 2018). The ENCODE data analysis revealed cMYC ChIP signals on both PTGES1 and 2 (Figure EV 4A-4B), which was confirmed in MDA-MB-436 cells (Fig 4C and 4D). However, we found that the binding of cMYC on PTGES1 promoter was unaltered by the miR-155 knockdown, suggesting alternative regulation on PTGES1. Further analysis in MDA-MB-436 and Hs578T breast cancer cells showed that the expression of cMYC restores the miR-155 knockdown-induced reduction in PTGES2 promoter activity(Fig 4E and 4F). This result indicates that cMYC contributes, at least in part, to the regulation of PTGES2 expression.

**Figure 4.**
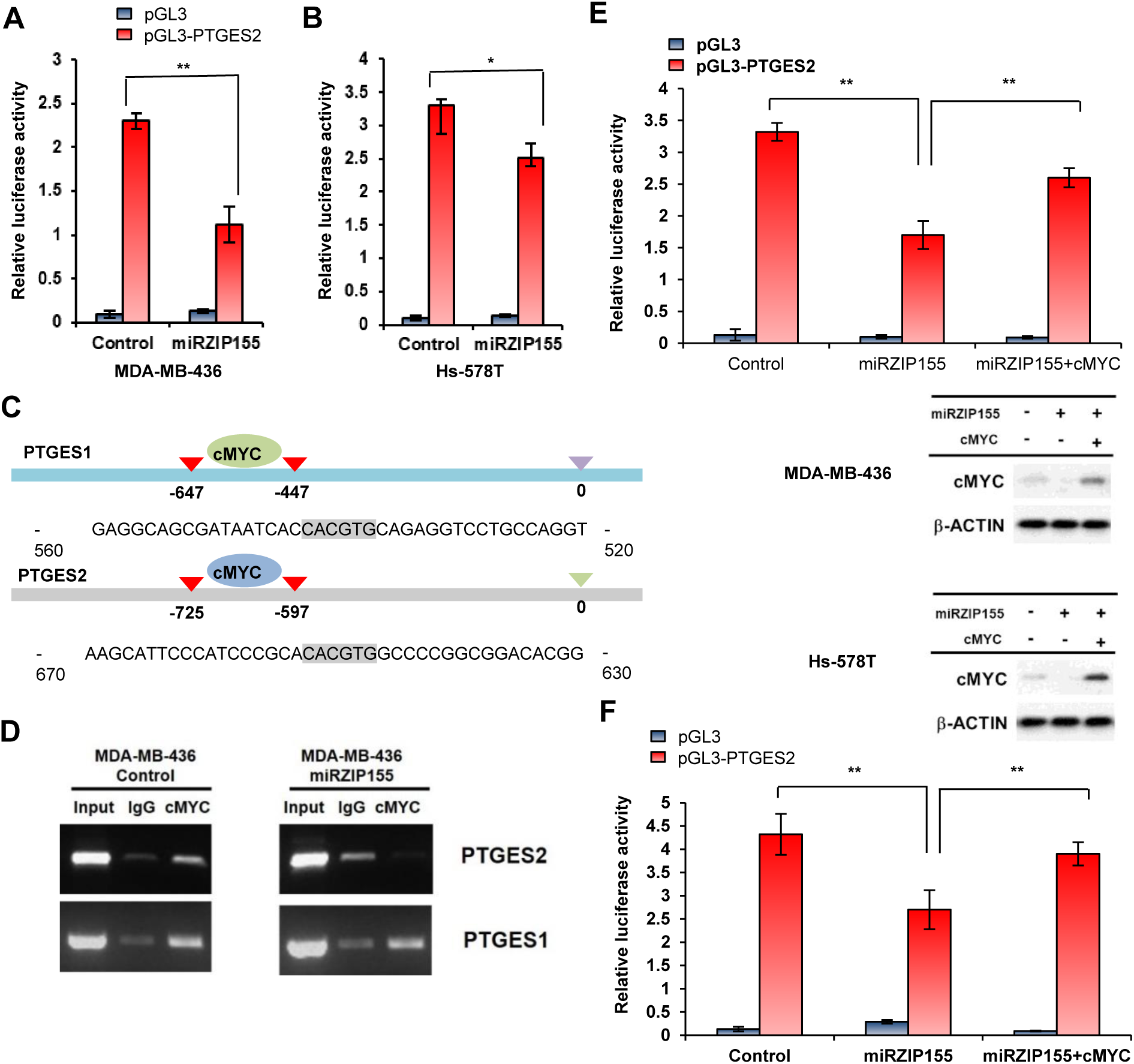
cMYC mediates the miR-155-driven PTGES2 transactivation. A and B. Results of the promoter reporter assay for PTGES2 in MDA-MB-436 (A) and Hs-578T cells (B) with (miRZIP155) or without (control) miR-155 knockdown. PGL3 empty vector was used as negative control. C. Schematic diagram showing the cMYC-binding site (E-box) on the PTGES1 and PTGES2 promoter. Numbers indicate nucleotide positions counted from the transcription start site. D. Chromatin immunoprecipitation result of cMYC in MDA-MB-436 control and knockdown (miRZIP155) cells. Promoters of PTGES1 and PTGES2 are tested. IgG was used as negative control. E and F. Results of the promoter reporter assay of PTGES2 in MDA-MB-436(E) and Hs-578T cells (F) with miR-155 knockdown (miRZIP) in combination with cMYC reconstitution (miRZIP155 + cMYC). Data on the two middle panels indicate the level of cMYC protein expressed in the reporter assay.

### PTGES1 expression is positively regulated by the miR-155-KLF4 axis

As the ChIP data for PTGES1 in Fig 4D suggested another mediator regulating PTGES1 expression by miR-155, we searched for such mediator. Promoter assay analysis for PTGES1 showed that miR-155 knockdown increases the promoter activity of PTGES1 (Fig 5A and 5B), implying a transcription activator regulated by miR-155. Among the candidates, KLF4 drew our attention as a recent report showed a positive correlation between KLF4 and PGE2 in M2 macrophages (Luan et al, 2015). Initial analysis of transcription factor binding predicted a site 184 nt upstream of TSS (Fig 5C). We confirmed KLF4 binding by ChIP assay and found that miR-155 knockdown eliminated the binding (Fig 5D right panel). Subsequent analysis revealed down-regulated KLF4 expression in both RNA (Fig 5E left) and protein (Fig 5E right) levels in miR-155 knockdown cells, indicating that KLF4 is positively regulated by miR-155. Subsequent promoter assay with KLF4 expression revealed that KLF4 can restore the miR-155 knockdown-induced reduction in promoter activity, supporting the idea that KLF4 mediates the regulation of PTGES1 expression by miR-155.

**Figure 5.**
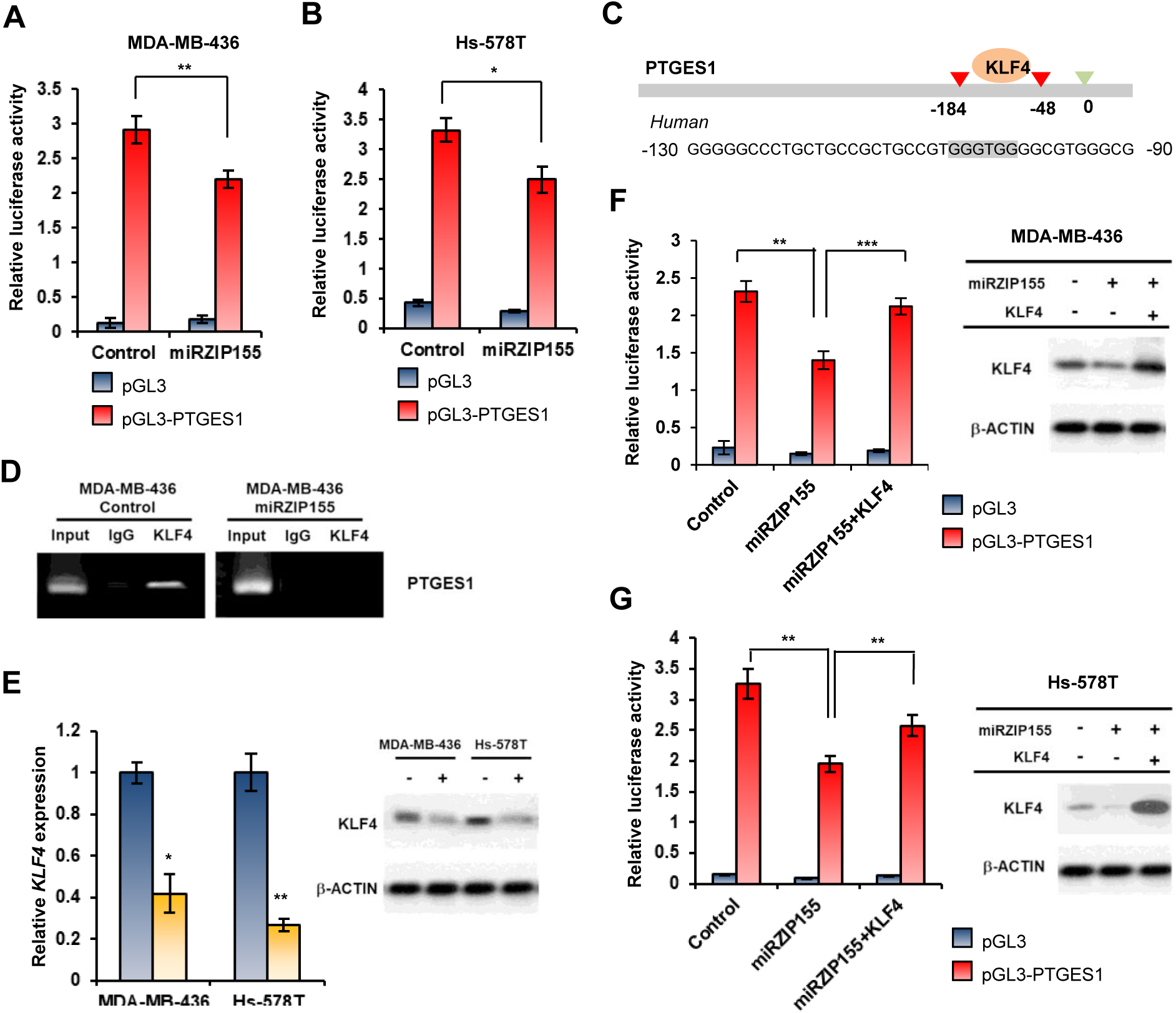
KLF4 mediates the miR-155-driven PTGES1 transactivation. A and B. Relative luciferase activity of the PTGES1 promoter reporter in MDA-MB-436(A) and Hs-578T(B); control (in blue) or miR-155 knockdown (in red) cells were tested. C. Schematic diagram showing the KLF4-binding site on the PTGES1 promoter. Numbers indicate nucleotide positions counted from the transcription start site. D. Chromatin immunoprecipitation result of KLF4 in MDA-MB-436 control and knockdown (miRZIP155) cells. E. Relative RNA expression of KLF4 in MDA-MB-436 and Hs-578T cells. Control (in blue) or miR-155 knockdown cells (miRZIP155, in yellow) were tested. Data on the right panel show KLF4 protein detected by Western blot. F and G. PTGES1 promoter reporter assay with KLF4 reconstitution, in MDA-MB-436 or in Hs-578T cells. The panels at the bottom show the KLF4 protein level measured by Western blot.

### Expression of miR-155, PTGES1/PTGES2, PTGDS, and PGE2/PGD2 confirms miR-155-dependent prostaglandin regulation in TNBC specimens

Our previous analysis showed that miR-155 is highly expressed in TNBC (Chang et al, 2011). Hence, we further examined if the role of mIR-155 in prostaglandin metabolism observed in breast cancer cells is also valid in TNBC specimens. First, we measured the level of PGE2/PGD2 in 20 TNBCs with various expression levels of miR-155 and observed a significant positive correlation between PGE2/PGD2 ratio and miR-155 (p=0.045; Fig 6A, raw data in Appendix Table S5). Moreover, when we divided the 20 tumors into miR-155-high and -low groups, we observed that miR-155-high group had higher PTGES1, PTGES2 and lower PTGDS expressions than the miR-155-low group (Fig 6B-6D). In contrast, there was no significant difference between the two groups in terms of the level of 15-PGDH (Fig 6E). The correlations in RNA level were further validated by Western blot analysis for the three enzymes as well as the suggested mediators (cMYC and KLF4; Fig 6F). Moreover, a correlation between miR-155 and PGE2/PGD2 level in the plasma of TNBC patients also supported our results (Fig 6G and 6H, patient information in Appendix Table S6 and raw data in Appendix Table S7). Collectively speaking, these results show that miR-155 regulates PGE2/PGD2 through PTGES1, PTGES2, and PTGDS. At this point, the molecular mechanism for the PTGDS regulation is not clear, but we speculate that a mode of action similar to the cMYC/KLF4-PTGES2/PTGES axis might be involved.

**Figure 6.**
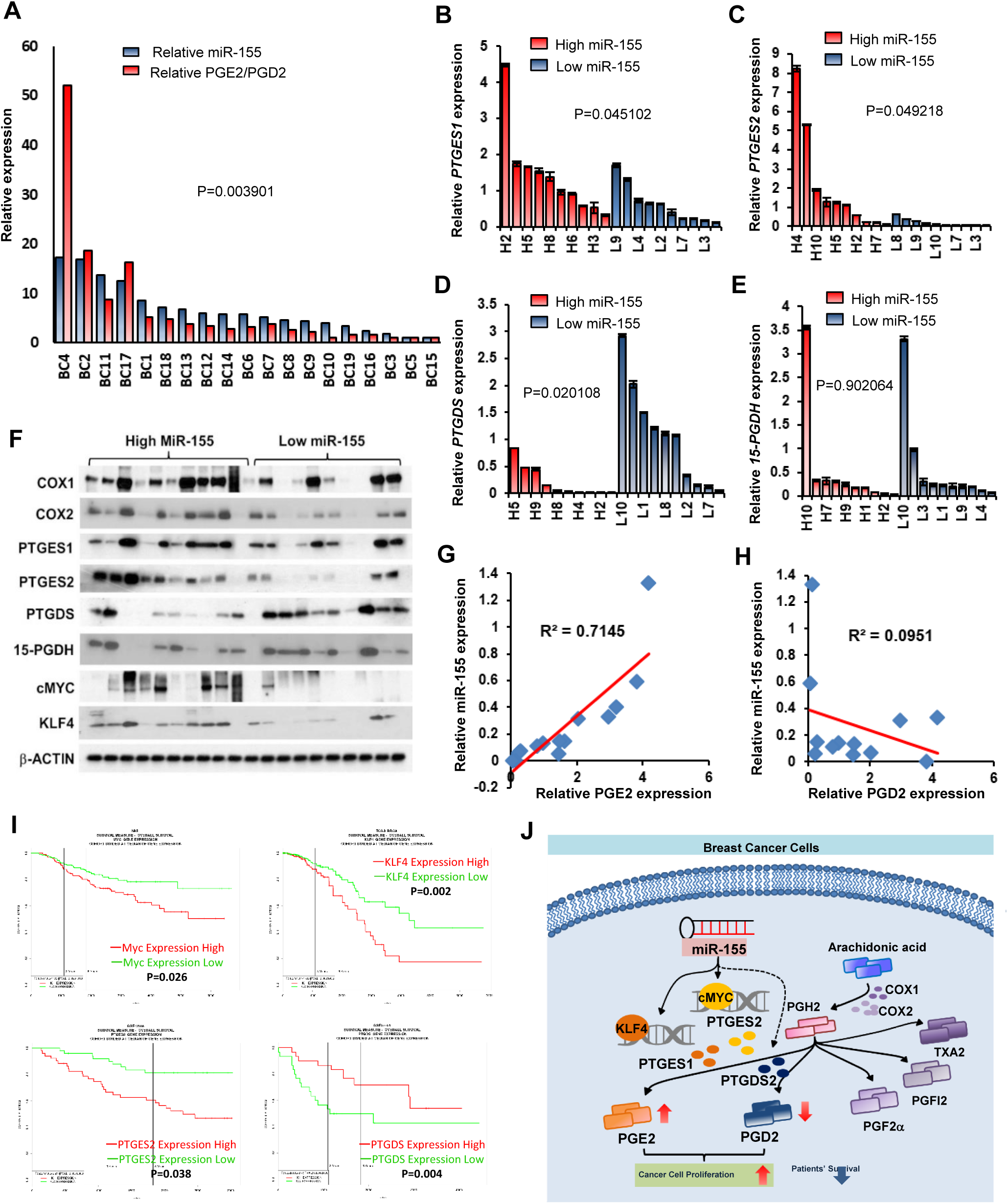
miR-155 positively correlates with PGE2/PGD2 ratio in tumor specimen and its target genes affect survival of breast cancer patients. A. Correlation analysis between miR-155 (in blue) and PGE2/PGD2 ratio (in red) in human TNBC specimens. p=0.045, B-E. Relative RNA expression of *PTGES1*(b), *PTGES2*(c), *PTGDS*(d) and *15-PGDH*(e) in miR-155-high (H1-H10, in red) and miR-155-low (L1-L10, in blue) TNBC samples. F. Western blot results of key metabolic enzymes and suggested novel regulators for PGE2/PGD2 metabolism in the miR-155-high and -low TNBC specimens. Results of COX1, COX2, PTGES1, PTGES2, PTGDS, 15-PGDH, cMYC, and KLF4 are shown. β-actin was used to ensure equal loading. G and H. Correlation analysis between miR-155 and PGE2(g) or PGD2(h) in the plasma samples of selected TNBC patients. I. Overall survival curves of breast cancer patients according to the levels of cMYC(upper left), KLF4(upper right), PTGES2(lower left) and PTGDS(lower right), using the ProgeneV2 webtool. J. Graphical abstract showing the molecular mechanism of PGD2/PGE2 production regulated by miR-155.

### miR-155 exerts its oncogenic potential by the regulation of PGE2 in breast cancer cells

Accumulating evidence indicate that PGE2 exerts oncogenic function in cancer via its receptor EP4 and its downstream signaling, including Ras-Raf-MEK-ERK and GSK3-beta/β-Catenin (Ho et al, 2015). In order to delineate the functional impact of miR-155-mediated PGE2/PGD2 metabolic shift, we selected two primary breast cancer cells and MCF7 with low miR-155 expression (Fig 3A and Figure EV 2B). We overexpressed miR-155 using lentivirus and treated them with siRNA of PTGES1 and PTGES2, to examine the contribution of those genes on miR-155 induced-oncogenicity. Cell proliferation assay results showed that miR-155 overexpression markedly increased the proliferation of the three breast cancer cells (Figure EV 5A-5C), which is in line with previous findings. Importantly, we observed that knockdown of PTGES1 or PTGES2 significantly reduced the miR-155-induced proliferation and that the combination of the two siRNAs nearly abolished the miR-155-induced proliferation in PDCL3 and MCF7 cells. We further tested the effect of PTGES1/2 knockdown in another experiment where cell proliferation was triggered by PGE2 treatment (Figure EV 5D-5F). These data demonstrate that miR-155 exerts its oncogenicity, at least in part, by increasing the PTGES1/2 level. Finally, we addressed if the expression of metabolic enzyme genes as well as their regulators identified in this study are associated with patient survival. Using PROGgeneV2 webtool (Goswami & Nakshatri, 2014), we found that higher expression of cMYC, KLF4, and PTGES2 was associated with poor survival whereas higher expression of PTGDS was associated with better survival in patients with breast cancer (Fig 6I). Taken altogether, our data demonstrate that miR-155-mediated PGE2/PGD2 regulation is an important factor for the oncogenicity and prognosis of breast cancer (Fig 6J).

## Discussion

Several reports have shown the impact of PGE2 in breast cancer, most of which have focused on the role of Cox1 and Cox2 that catalyzes PGH2 production from arachidonic acids (Chang et al, 2005; Howe, 2007; Subbaramaiah et al, 2012; Xu et al, 2014). Initial observation on the association between miR-155 and Cox expression came from the studies on immunological diseases such as asthma (Comer et al., 2015). Later studies reported elevated levels of miR-155 and Cox2 in various cancers including colorectal cancer (Comer, 2015). In this report, we demonstrated a novel function of miR-155 in prostaglandin metabolism via the regulation of PTGES1/2 and PTGDS, which are directly responsible for the synthesis of PGE2 and PGD2, respectively. Even though PTGES1 and PTGES2 share the same function, their localization is different: the former resides in the perinuclear zone and belongs to the MAPEG (for membrane-associated proteins involved in eicosanoid and GSH metabolism) family, whereas the latter is a Golgi membrane-associated protein, and the proteolytic removal of the N-terminal hydrophobic domain leads to the formation of a mature cytosolic enzyme (Murakami & Kudo, 2006). The finding that miR-155 controls both enzymes suggests that it has a broad range of functional outcomes. Related to this, siRNA treatment results in the Supplementary Fig 6A-6C show that the knockdown of PTGES1/2 does not fully restore the miR-155-induced proliferation, suggesting that other mechanisms of miR-155 might have induced proliferation as well as the effect of PTGDS suppressed by miR-155.

The regulation of PTGES1 has been extensively studied (Ramanan & Doble, 2017), and in our study, we identified KLF4 as a novel regulator for the PTGES1. A previous report showed that KLF5 is involved in this regulation (Xia et al, 2013). KLF family consists of 17 members that bind to GC-rich promoter regions of the genes (consensus (G/A)(G/A)GG(C/T)G(C/T)) involved in proliferation, differentiation, and apoptosis (Ghaleb et al, 2005; Kaczynski et al, 2003). We initially investigated KLF4 as a positively correlated gene of Fos transcription factor (Figure EV 6A), which is known to be up-regulated by miR-155 (Ning et al, 2017). Because miRNA generally suppresses its direct target genes, their direct targets generally show inverse correlation with their expression. However, as KLF4 showed a positive correlation with miR-155 expression (Fig 5E), we speculated that KLF4 is an indirect target of miR-155. Indeed, UTR analysis of KLF4 did not show any miR-155 binding sites (Appendix Figure S1). A recent report demonstrated that Fos is up-regulated by miR-155 in MCF7 cells (Martin et al, 2014), which is in line with our results. Interestingly, miR-155 is also reported to be positively regulated by AP-1 (composed of Fos and Jun transcription factors) in B-cells (Yin et al, 2008), suggesting a possible feedback-loop between Fos and miR-155 expression.

As for PTGES2, we observed miR-155-dependent interaction with cMYC, which was previously reported to be upregulated by miR-155 via Foxo3a (Kim et al., 2018). Thus, both PTGES1 and 2 are indirectly up-regulated by miR-155. On the other hand, the regulation of PTGDS is not fully known. In adipocytes, one report showed that the mineralocorticoid receptor (MR) targets PTGDS (Urbanet et al, 2015) and another showed that PTGDS is associated with estrogen level (Lim et al, 2015). The regulation of PTGDS by miR-155 remains unclear as well. There is no predicted miR-155 binding site in the mRNA of PTGDS. There are known binding sites of six transcription factors on the PTGDS promoter, including HIC, ZFHX2, GLIS1, HLF, ZIC2, and EGR2. However, none of them are known to be regulated by miR-155 or have Ago-CLIP seq signal altered by miR-155 depletion (Loeb et al, 2012). Therefore, we speculate that other mechanisms such as epigenetic regulation or involvement of novel transcription factors are responsible for this regulation.

Our study suggests that the abnormal PGE2 production in cancer cell may be reversed by inhibition of miR-155 (Fig 2A). Most of the trials on reducing PGE2 level have been focused on the inhibition of Cox1 or Cox2, but the inhibitors showed cardiac toxicity (Dowd et al, 2001), thereby preventing further development of these inhibitors in clinical trials. As the antagomiR of miR-155 (MRG-106) successfully underwent phase 1 clinical trial for lymphoid malignancy (Seto et al, 2018), our data suggest an alternative way of inhibiting PGE2 in breast cancer by miR-155 inhibition. Additionally, a recent study showed that the Cox2/PTGES1/PGE2 pathway increases PDL1 expression in tumor-associated macrophages and MDSCs (Majumder et al, 2018), suggesting that miR-155 can aid tumor immunosurveillance by stimulating PGE2 expression. Further studies are needed to clarify such regulation and to expand the use of anti-miR-155 for cancer immunotherapy.

## Materials and Methods

### Cell culture and transfection

The miR-155-deficient breast cancer mouse model was generated as previously described (Kim et al., 2016). miR-155 heterozygous (miR-155 ^KO/+^) or knockout (miR-155 ^KO/KO^) primary cancer cells were isolated from tumor tissues using a modified MEC (mammary epithelial cell) isolation procedure as previously described (Chang et al., 2011). The mouse primary cells—MCF7, Hs-578T, MDA-MB-436, HeLa, and 293TN (System Biosciences, SBI, Mountain View, CA, USA)—were maintained in DMEM containing 10% FBS and 1% antibiotics. Human primary breast cancer cells obtained from patients with triple-negative breast cancer (TNBC) (Asan Medical Center, IRB No. 2013-0939) were previously described (Jung et al, 2018). Depending on the level of miR-155, six of the primary cells were grouped as patient-derived cells (PDCs) H1-H3 and L1-L3. The human primary cells were cultured in RPMI1640 medium with 10% FBS, hEGF (200 ng/ml), hydrocortisol (10 mg/ml), transferrin (10 mg/ml), and 1% penicillin/streptomycin. All plasmids and siRNAs were transfected into different cells using lipofectamine 2000 (Invitrogen) according to the manufacturer’s instructions. Small interfering RNAs (siRNAs) against human PTGES1 and PTGES2 were purchased from Genolution Inc. (Seoul, Republic of Korea). The siRNA sequences of *PTGES1* and *PTGES2* were as follows: *PTGES1*, 5′-UUGUUGGCUCACAGUAGUGG-3′; *PTGES2*, 5′-UGAUGAUCCAAGAGCUCUUGCC-3′. PDCL2,3 and MCF7 cells were transfected with either 200 nmol/L of target siRNAs (*PTGES1* or *PTGES2*) or scramble siRNA using lipofectamine 2000.

### Enzyme-linked immunosorbent assay (ELISA)

Blood samples were obtained from patients with triple-negative breast cancer (TNBC) (Asan Medical Center, IRB No.2013-0939). The plasma concentrations of PGE2 and PGD2 were determined using ELISA kits from ENZO Life Sciences (Farmingdale, New York, NY, USA) and CUSABIO Biotechnology (Wu Han, China), respectively. An aliquot of 10 µl of serum added to 90 µl of diluent (0.1% bovine serum albumin, 0.05% Tween 20, 10 mM Tris and 150 mM sodium chloride, pH 7.3) was used for analysis.

### LC-MS/MS analysis

A liquid chromatography-tandem mass spectrometry (LC-MS/MS) system was used to measure metabolites according to a previously described method (Kim et al., 2015).

### Quantitative RT-PCR (qRT-PCR)

Total RNA extraction was performed using TRizol (Invitrogen) according to the manufacturer’s instruction. The primer sequences are shown in Supplementary Table 8. The mRNA expression levels of major eicosanoids related genes (*PTGES1, PTGES2, PTGDS, 15-PGDH*) and *KLF4* genes were measured by real-time RT-PCR according to the manufacturer’s protocol. Relative expression value was normalized to *RPL13a* and calculated by using the 2-(ΔΔCt) method. Quantitative measurement of miR-155 was performed according to the previously described method (Chang et al., 2011). The expression was normalized with small nuclear RNA U6 (RNU6).

### Antibody and Western blot analysis

Preparation of total cell lysates and Western blot analysis were performed as previously described (Jung et al., 2018). A total of 10–50 µg of protein was used per lane. The blot was probed with anti-COX1 (1:1000, Cell Signaling Technology, Danvers, MA, USA), anti-COX2 (1:1000, Cell Signaling), anti-PTGES1 (1:500; Santa Cruz Biotechnology, Santa Cruz, CA, USA), anti-PTGES2 (1:1000, Santa Cruz Biotechnology), PTGES2 (1:1000, Santa Cruz Biotechnology), anti-PTGDS (1:1000, Santa Cruz Biotechnology), anti-15-PGDH (1:1000, Santa Cruz Biotechnology), anti-KLF4 (1:1000, Cell Signaling), anti-cMYC (1:1000, Cell Signaling) and anti-β-actin (1:1000, Santa Cruz Biotechnology) antibodies. The relative densities of bands were analyzed with the NIH Image J 1.47v software.

### miR-155 knockdown or overexpression

Lentiviral vector system from SBI (Mountain View, CA, USA) was used for miR-155 inhibition (miRZIP155). The lentiviral production was performed according to the manufacturer’s instructions. Control or miRZIP155-containing lentivirus particles were infected into MDA-MB-436 and Hs-578T. After 24 hours, cells were selected by Puromycin (1 mg/ml; Sigma) for 7 days. For miR-155 overexpression, a lentivirus was made using a miR-155 overexpression vector (miRH155 or control) according to the same procedure as miRZIP155 virus. MCF7 and human primary breast cancer cells (PDCL1-L3) were infected with miRH155 lentiviral vectors for 24 h.

### Chromatin immunoprecipitation (ChIP) assay

ChIP was performed according to a protocol (http://cshprotocols.cshlp.org/content/2009/9/pdbprot5279. full) with minor modifications. Briefly, MDA-MB-436 cells infected with control or miRZIP155 lentiviral vectors were cross-linked with 1% formaldehyde for 10 minutes at RT. The cells were then collected, lysed, sonicated, and incubated for overnight with a C/EBP-β antibody. PCR was used to detect ChIP signal with the primers listed in Appendix Table S8.

### Reporter construction and luciferase assay

For reporter construction, the promoter regions of PTGES1 and PTGES2 were amplified (primer sequences in Appendix Table S8) and introduced into the pGL3 Enhancer vector (Promega) via *Spe*I and *Hin*dIII sites. MDA-MB-436 and Hs-578T cells (infected with control or miRZIP155 lentiviral vectors) were transfected with reporter vectors (pGL3-PTGES1 or pGL3-PTGES2 or Empty vector) with or without overexpression vector (pcDNA3-cMYC or pEIGW-human KLF4, gift from Dr. Han Seok Choi, University of Ulsan College of Medicine, Seoul, Republic of Korea). After 48 hours, luciferase activity was measured using Dual-Luciferase Reporter Assay System (Promega Corp, WI, USA) and Victor Luminometer (Perkin-Elmer, MA, USA).

### alamarBlue^®^ cell viability assay

To test whether the inhibition of *PTGES1* or *PTGES2* using siRNAs can reverse the increase in the growth of cancer cell triggered by miR-155 or PGE2, we performed two methods using lentiviral vector encoding miR-155 (miRH155) and PGE2. The cells infected with miRH155 were transfected with siPTGES1 or siPTGES2 in a 96 well plate. After 48 hours, 1/10^th^ volume of alamarBlue^®^ reagent (Invitrogen) was directly added to the culture media. The cells were incubated for additional 4 hours to assay viability, which was detected by fluorescence measurements using a microplate fluorescence spectrophotometer (GenTeks Biosciences, Inc., San-Chong, Taipei). For induction of proliferation by PGE2, the cells were pre-treated with 100 mM PGE2 for 4 days and transfected with siPTGES1 or siPTGES2 plus PGE2. The procedure is same as mentioned above.

### Human TNBC sample analysis

Human cancer samples were obtained from patients diagnosed with triple negative breast cancer (TNBC) (Asan Medical Center, IRB No.2013-0939). We used 71 formalin-fixed, paraffin-embedded (FFPE) tissues of TNBC samples as previously described (Kim et al., 2018). Briefly, total miRNAs were extracted from 8-µm thick FFPE tissues by using a miRNeasy FFPE kit (Qiagen). Among them, we selected 15 samples from each group, based on the frozen tissue availability at our Bio Resource Center.

### Statistical analysis

All data are reported as mean ± standard error. Comparisons between two groups were performed using the Student’s *t*-test and among multiple groups by ANOVA. *P* values < 0.05 were statistically significant.

## Acknowledgments

This research was supported by the Basic Science Research Program of the National Research Foundation Korea (NRF), funded by the Ministry of Education (grant number: KNRF-2018080504 and 2014R1A1A2055907)

2017

## Author contributions

Sinae Kim and Eun Sung Lee performed main experiments and primary data analyses. Eun ji Lee and Jae Yun Jung provided technical supports in animal and cellular study. Sae Byul Lee, Jisun Kim, Hee Jung Kim and Jong Won Lee provided tumor specimen and clinical information of tumors, under the control of IRB. Hee Jin Lee and Gyungyub Gong performed pathological analysis of tumor and helped on miRNA quantitation from FFPE. Byung Ho Son and Sei Hyun Ahn contributed on study design and financial support. Suhwan Chang conceived idea, supervised the research progress, designed analysis strategies, and wrote the manuscript. All the authors have read and approved the final manuscript.

## Expanded View Figure Legends

**Figure EV1**. Profile of eicosanoids in mammary breast tumor cell lines (Lav670 and Lav714) and its subclone with stable expression of miR-155 knockdown vector, marked as miRZIP155. Relative fold change in metabolite level is shown for each of the eicosanoids detected.

**Figure EV2**. A and B. miR-155 expression measured in MDA-MB-436 and Hs-578T cells after knockdown (A) or in MCF7 cells after overexpression (B). C. Eicosanoid profiling in human breast cancer cells with miR-155 depletion or overexpression. In MDA-MB-436 and Hs-578T cells, miR-155 was knocked-down (miRZIP155). In MCF7 cells, miR-155 was overexpressed (miRH155).

**Figure EV3**. miRNA binding prediction for PTGES UTR (A) or PTGES2 UTR (B). The images were taken from the TargetScanHuman 7.1 webtool. (http://www.targetscan.org/vert_71/)

**Figure EV4**. Transcription factor binding prediction on the PTGES(A) or PTGES2(B) promoter. The cMYC binding predictions were marked with red boxes. The images were obtained from the UCSC genome browser (https://genome.ucsc.edu/)

**Figure EV5**. Transient knockdown of PTGES/PTGES2 restores miR-155 or PGE2-induced cellular proliferation.

A-C. Proliferation of miR-155-low PDC (PDCL2 and PDCL3, A and B) and MCF7(C) after miR-155 overexpression (miRH155) or treatment in combination with siRNAs for PTGES1/PTGES2.

D-F. Proliferation of miR-155-low PDC (PDCL2 and PDCL3, D and E) and MCF7(F) after treatment with PGE2 or in combination with siRNAs for PTGES1/PTGES2.

